# *Drosophila melanogaster* lactate dehydrogenase deficiency recapitulates the exercise intolerance of human glycogen storage disease type XI

**DOI:** 10.64898/2026.07.09.736989

**Authors:** Madhulika Rai, Shefali A. Shefali, Jason P. Tourigny, Minseo Kim, Travis Nemkov, Angelo D’Alessandro, Jason M. Tennessen

## Abstract

Lactate dehydrogenase A (LDHA) is a key glycolytic enzyme that commonly exhibits altered expression in human diseases such as cancers and neurodegeneration, making it a valuable disease biomarker and putative therapeutic target. However, any treatment targeting LDHA will also disrupt normal metabolism, underscoring the need to investigate physiological consequences of inhibiting this enzyme. We previously established the fruit fly *Drosophila melanogaster* as a genetic model for studying LDH function in the context of growth, metabolism, and development. Here we expand upon those studies by investigating a serendipitous observation that *Ldh* mutant larvae exhibit diet-dependent lethality. Using a multiomic approach, we discovered this diet-dependent phenotype is independent of nutritional composition. Instead, *Ldh* mutant larvae are exercise intolerant and display reduced mobility, rendering mutant larvae sensitive to food consistency. Moreover, tissue-specific analysis reveals that LDH activity within muscle and peripheral glia are essential for larval viability raised on solid food. Intriguingly, these phenotypes mirror the pathophysiology of LDHA deficiency (Glycogen Storage Disease Type XI; GSD Type XI) in humans, where mild symptoms are exacerbated by physical exertion and environmental stress. Together, our findings further highlight the value of using *Drosophila* to explore the developmental and physiological consequences of Ldh inhibition.

**SUMMARY STATEMENT:** Loss of lactate dehydrogenase in *Drosophila* causes exercise intolerance and diet-dependent lethality, revealing conditional phenotypes that mirror those associated with human LDH-A deficiency.

## INTRODUCTION

Lactate dehydrogenase (LDH) catalyzes the reversible conversion of pyruvate and lactate, a reaction essential for maintaining cellular redox balance and sustaining glycolytic flux (Rabinowitz and Enerback, 2020, Lin et al., 2022, Farhana and Lappin, 2026). In vertebrates, the LDHA isoform plays a particularly important role in energy-demanding tissues such as skeletal muscle and brain, where it supports rapid ATP production under both aerobic and anaerobic conditions (Bartoloni et al., 2024, Kumar et al., 2025, Kim et al., 2025). However, Ldh functions in cellular processes that extend beyond maintaining redox balance, with recent studies demonstrating roles for both Ldh and lactate in signal transduction, chromatin modification, and gene regulation (Ferguson et al., 2018, Brooks, 2018, Zhang et al., 2019, Liberti and Locasale, 2020, Liu et al., 2023). Consequently, LDHA has emerged as a key regulator of biological programs ranging from embryonic development to immune cell differentiation and neuronal function (Valvona et al., 2016, Peng et al., 2016, Chen et al., 2023, Seth et al., 2017, Dai et al., 2023, Tendulkar et al., 2025, Yao et al., 2023).

Consistent with its broad functional reach, LDH is frequently implicated in a variety of disease states. In cancer, elevated LDHA expression supports the metabolic demands of rapidly proliferating cells and contributes to immune evasion and tumor progression (Brand et al., 2016, Wen et al., 2025, Ma et al., 2025). Inflammatory and autoimmune conditions often involve altered lactate signaling, which can skew macrophage polarization and disrupt tissue homeostasis (Fang et al., 2024, Llibre et al., 2025, Liu et al., 2025). In the nervous system, Ldh dysfunction has been linked to impaired glial support and motor neuron degeneration, contributing to diseases such as amyotrophic lateral sclerosis (Tendulkar et al., 2026, Bloom et al., 2022). Similarly, cardiovascular and metabolic disorders also show evidence of LDH dysregulation, where lactate accumulation and redox imbalance exacerbate tissue damage (Sanchez et al., 2021, Zhu et al., 2022, Xiao et al., 2023). These diverse roles position Ldh not only as a metabolic enzyme but also as a central player in disease pathogenesis.

While LDHA inhibition has been repeatedly proposed as a promising therapeutic strategy, the systemic importance of this enzyme raises concerns about unintended side effects. A window into these risks is provided by Glycogen Storage Disease Type XI (GSD-XI), a rare autosomal recessive disorder caused by mutations in the *Ldha* gene (Kanno et al., 1980). Individuals with GSD-XI typically present with exercise intolerance, muscle pain and stiffness, and recurrent episodes of exertion-induced rhabdomyolysis, often accompanied by elevated serum creatine kinase levels and myoglobinuria. Some patients also develop characteristic psoriasis-like skin lesions, indicating that the consequences of LDHA deficiency extend beyond skeletal muscle (Serrano-Lorenzo et al., 2022, Ozen, 2007, Kanno et al., 1988). Collectively, these clinical manifestations underscore the essential role of LDHA in maintaining physiological homeostasis and illustrate how impaired LDHA activity can compromise the ability to respond to metabolic stress, highlighting potential liabilities associated with long-term therapeutic inhibition of this enzyme.

Since Ldh is highly conserved and lactate metabolism plays a central role in cellular physiology, Ldh function has been studied across a diverse array of organisms. Among these, the fruit fly *Drosophila melanogaster* stands out as an ideal system for investigating LDH inhibition. Under most conditions, flies express a single LDH enzyme whose core functions are conserved (Abu-Shumays and Fristrom, 1997). This simplicity contrasts with vertebrates, where multiple LDH isoforms complicate genetic analysis. Moreover, unlike mice, where *Ldha* mutations are embryonic lethal (Merkle et al., 1992), Ldh-deficient flies are viable and exhibit decreased lactate levels and excess glycogen accumulation (Li et al., 2019), thus mirroring the relatively mild phenotypes associated with GSD Type XI and providing a straightforward system for studying Ldh function. Additionally, the link between altered Ldh expression and disease progression is conserved in flies, with recent studies demonstrating roles for *Drosophila* Ldh in tumor growth, neurodegeneration, and aging, supporting the relevance of using flies to study Ldh in disease models (Wang et al., 2016, Herranz and Cohen, 2017, Long et al., 2020, Frame et al., 2023, Park et al., 2025). Finally, because Ldh expression is tightly coordinated with growth (Tennessen et al., 2011, Rai et al., 2024, Rechsteiner, 1970), *Drosophila* provides a powerful platform for uncovering how lactate metabolism influences cell growth, proliferation, and differentiation.

In this study, we extend these parallels between LDH deficiency in *Drosophila* and human GSD-XI by describing a diet-dependent lethal phenotype. Using metabolomic and transcriptomic profiling, we show that this lethality is not driven by nutrient composition or hypoxic microenvironments within the food but is instead associated with muscle dysfunction. In line with these findings, we demonstrate that survival depends on the physical consistency of the diet rather than its nutritional content, suggesting that physical exertion contributes to the lethal phenotype. Indeed, further analysis reveals that loss of Ldh activity restricts larval mobility and renders animals exercise intolerant. Finally, tissue-specific knockdown experiments identify muscle and peripheral glia as key sites of Ldh activity for viability on solid food. Together, these observations highlight *Drosophila* as a powerful model for both exploring the role of LDH promoting developmental growth and modeling the consequences of systemically inhibiting the activity of this highly conserved enzyme.

## RESULTS

### *Ldh* mutant larval viability is sensitive to culture conditions

We previously reported that *Drosophila Ldh* mutants exhibit mild developmental delays and partial lethality, with approximately 50% of larvae surviving to adulthood when raised on molasses agar plates supplemented with yeast paste (M+Y). However, while maintaining *Ldh*^−^/TM3 balanced stocks on standard Bloomington *Drosophila* Stock Center food (BDSC-F), we observed a complete absence of homozygous *Ldh* mutant pupae in the stock vials, suggesting that culture conditions strongly influence larval viability. To explore this possibility, we compared survival on different diets and found that while more than 50% of *Ldh* mutant larvae pupated on M+Y plates (Figure 1A; consistent with Li et al., 2019), nearly all larvae died when reared on BDSC-F (Figure 1A, S1A; see methods). These findings indicate that *Ldh* mutants are sensitive to culture conditions and suggest that environmental factors play a critical role in their viability.

**Figure 1.**
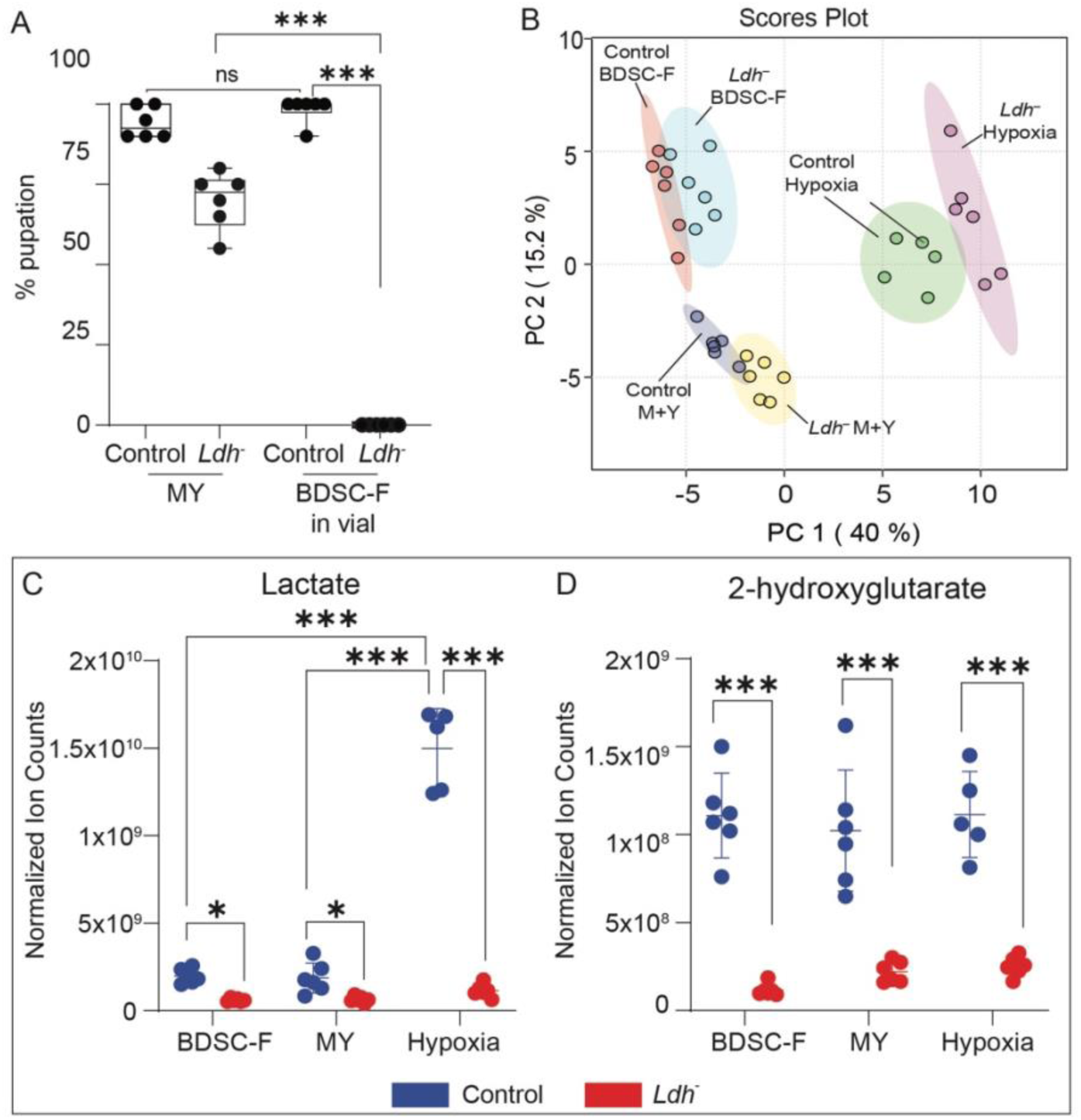
*Ldh* mutants exhibit diet-dependent lethality independent of global metabolic state. (A) Percent pupation of control (*Ldh^17/+^*) and *Ldh* mutant (*Ldh^17/16^*) larvae reared on M+Y plates or BDSC-F in vials. More than 50% of *Ldh* mutants pupariate on M+Y while nearly all larvae raised on BDSC-F die prior to the onset of metamorphosis. n ≥ 3 biological replicates with 10 larvae per replicate. Statistical analysis was performed using repeated-measures one-way ANOVA with Holm–Sidak correction (\*\*\**p* < 0.0001). (B) Principal component analysis (PCA) of metabolomic profiles. Samples segregate primarily by environmental condition, with hypoxia-exposed larvae separating from normoxic samples along PC1, and diet (M+Y vs. BDSC-F cubes) contributing to separation along PC2. Genotype also shows a shift along PC1 to a lesser extent. (C–D) Relative abundance of (C) lactate and (D) 2-hydroxyglutarate (2HG) in control (blue) and *Ldh* mutant (red) larvae raised on M+Y, BDSC-F cubes, or M+Y under hypoxia. *Ldh* mutants exhibit reduced levels of both metabolites across conditions, while hypoxia specifically elevates lactate in controls. Metabolite levels are comparable between diets under normoxic conditions. n ≥ 6 biological replicates. Statistical analysis was performed using two-way ANOVA with Holm–Sidak correction (\**p* < 0.05, \*\*\**p* < 0.0001).

### Diet-dependent Ldh mutant lethality is independent of nutrient composition

Given the central role of Ldh in maintaining cellular redox balance, we hypothesized that components of BDSC-F induce metabolic stress that is normally buffered by Ldh activity. To test this, we performed UHPLC-based metabolomic profiling of control and *Ldh* mutant larvae raised on either M+Y plates or BDSC-F (Figure S1A; Table S1). As a positive control, we also analyzed larvae raised on M+Y and exposed to hypoxia – a stress condition in which Ldh function is known to support metabolic adaptation. As an initial analysis of these data, we examined the abundance of lactate and 2-hydroxyglutarate (2HG), two metabolites synthesized by Ldh (Li et al., 2017). As expected, both metabolites were significantly reduced in *Ldh* mutants across all conditions and lactate levels were specifically elevated in hypoxia-exposed controls (Figure 1C–D). Importantly, control larvae raised on BDSC-F exhibited lactate and 2HG levels comparable to those observed in controls raised on M+Y (Figure 1C-D), indicating that these diets do not differentially alter steady-state levels of Ldh-derived metabolites.

To assess global metabolic differences, we performed principal component analysis (PCA) across 260 metabolites. This analysis revealed that samples primarily segregated by environmental condition rather than genotype – hypoxia-exposed larvae separated from normoxic samples along the PC1 axis, while M+Y- and BDSC-F-fed larvae separated along PC2 (Figure 1B). Notably, control and *Ldh* mutant samples within the same condition clustered more closely than identical genotypes across different conditions, indicating that diet and oxygen availability exert a stronger influence on the metabolome than loss of Ldh activity. However, we would note that the effect of Ldh loss also presents as a shift along the PC1 hypoxic axis, raising the possibility that loss of Ldh has a lesser, but similar effect as hypoxia.

We next asked whether the dietary condition that causes lethality in *Ldh* mutants was associated with greater metabolic disruption. Contrary to this expectation, *Ldh* mutants exhibited more extensive metabolic alterations when raised on M+Y plates than on BDSC-F (Figure 2A–B), despite exhibiting increased lethality on BDSC-F. Specifically, 38 metabolites were significantly altered in M+Y-fed mutants (fold change ≥ 2 and adjusted p-value ≤ 0.1; Figure 2B, Table S2), compared to only 7 metabolites in mutants fed BDSC-F (Figure 2A, Table S3). Together, these findings indicate that the diet-dependent lethality of *Ldh* mutants cannot be readily explained by differences in nutrient composition or overall metabolic state, suggesting that other features of the feeding environment likely underlie this phenotype.

**Figure 2.**
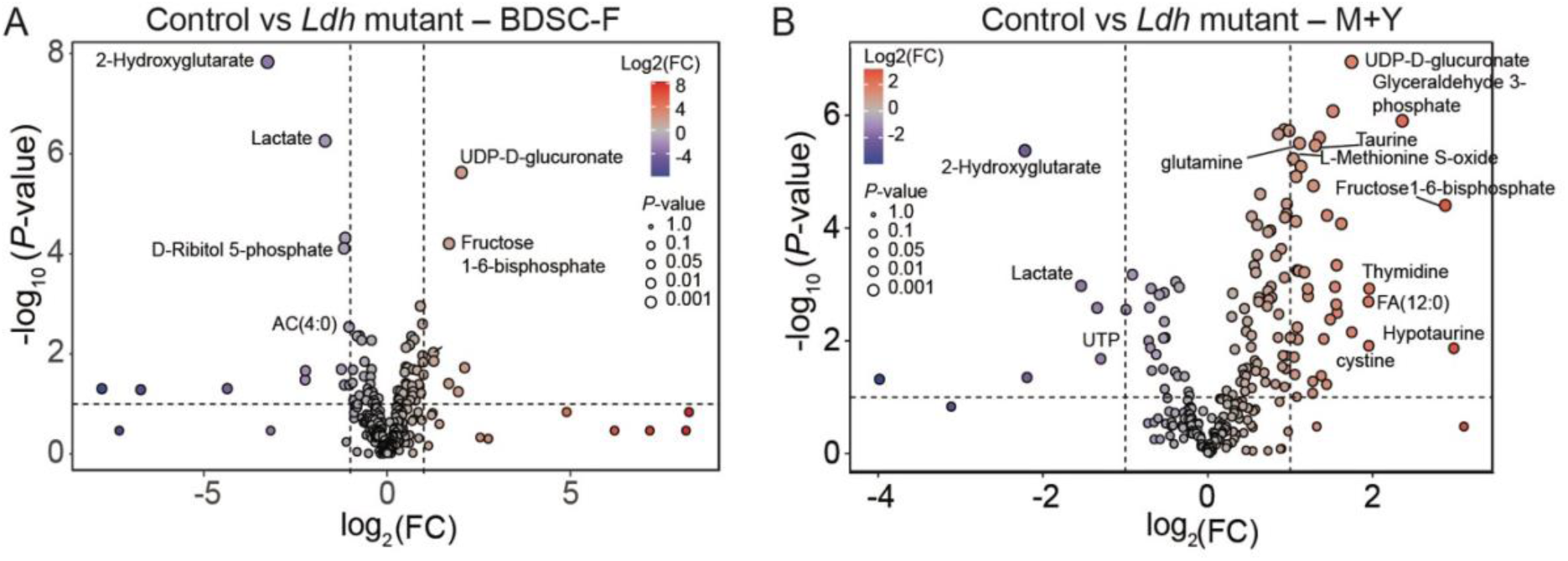
Diet-dependent *Ldh* mutant lethality does not correlate with nutrient composition. (A–B) Volcano plots showing metabolite abundance changes between control (*Ldh*^17^/+) and *Ldh* mutant (*Ldh*^17^/*Ldh*^16^) larvae raised on (A) BDSC-F cubes or (B) M+Y food. Metabolomic analyses were performed using six biological replicates per condition, with each replicate consisting of 20 larvae. Statistical analyses were conducted using MetaboAnalyst 6.0 as described in the Methods.

To further examine potential causes of the diet-dependent lethal phenotype, we performed RNA-seq on control and *Ldh* mutant larvae raised on M+Y or solid BDSC-F (Table S4). Consistent with the metabolomic analyses, *Ldh* mutants exhibited more extensive transcriptional changes on M+Y (n = 711 DEGs) than on BDSC-F (n = 402), despite the increased lethality observed on BDSC-F. Moreover, the majority of DEGs identified in mutants fed BDSC-F (310 of 402) overlapped with those altered in mutants raised on M+Y (Figure 3A–B), indicating that loss of Ldh activity elicits a similar transcriptional response when exposed to either dietary condition.

**Figure 3.**
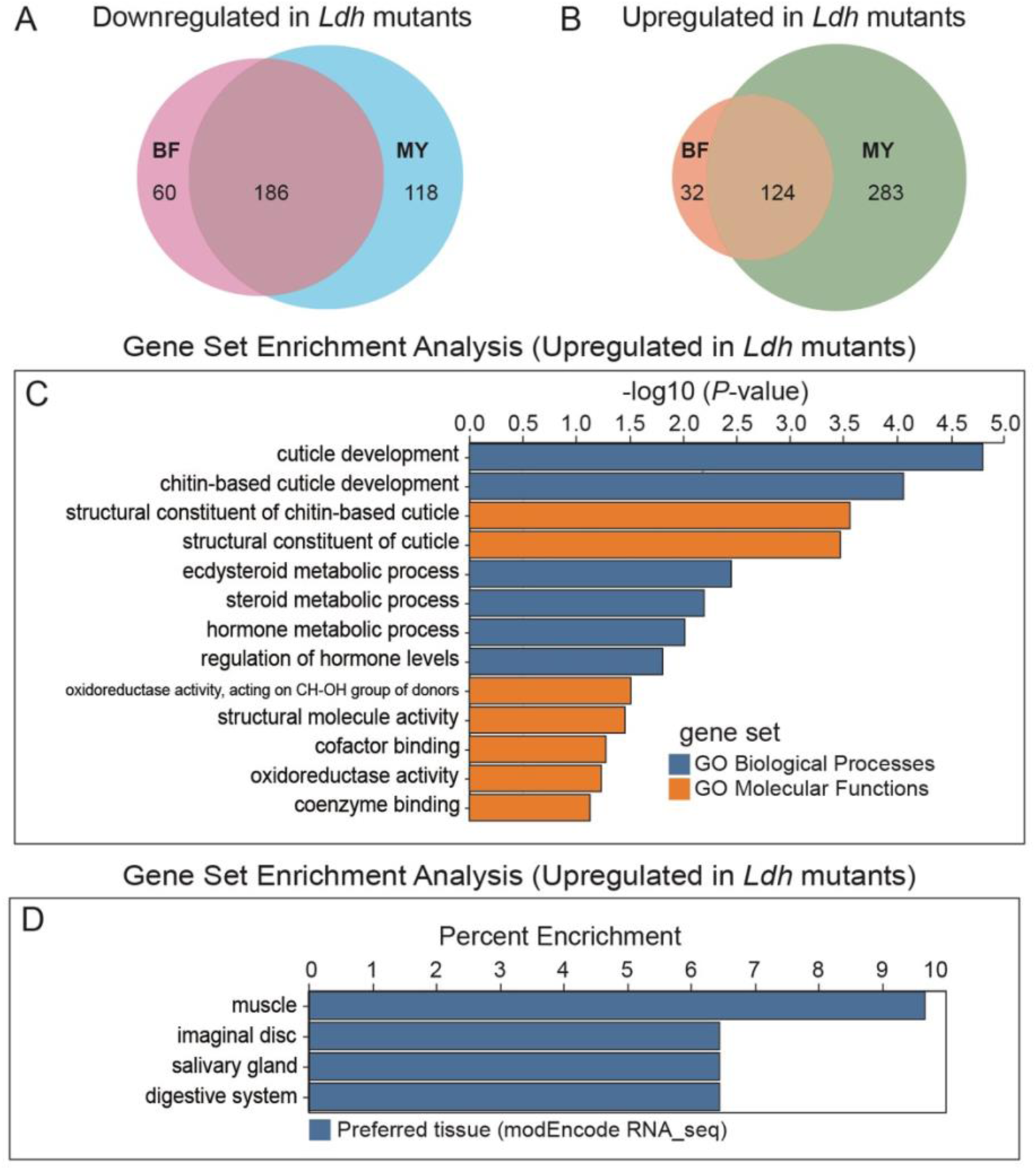
*Ldh* mutants raised on BDSC-F exhibit enrichment of cuticle- and muscle-associated gene expression programs. RNA-seq was used to compare gene expression between *Ldh* mutant (*Ldh*^17^/*Ldh*^16^) and control (*Ldh*^17^/+) L2 larvae. (A–B) Venn diagrams showing the overlap among genes that were (A) downregulated or (B) upregulated in *Ldh* mutants raised on M+Y or BDSC-F cubes relative to controls. Most genes differentially expressed in BDSC food overlap with those in Molasses + Yeast. (C) Gene ontology enrichment analysis of the 32 genes uniquely upregulated in *Ldh* mutants raised on BDSC-F cubes. Genes were analyzed for enrichment of GO Biological Process and GO Molecular Function categories using PANGEA, revealing significant enrichment of cuticle- and chitin-associated structural functions. (D) Preferred tissue enrichment analysis of these genes revealed significant enrichment for muscle-associated expression. RNA-seq analyses were performed using three biological replicates per condition, with each replicate consisting of 20 larvae.

To identify features uniquely associated with the lethal condition, we focused on genes specifically altered in *Ldh* mutants raised on BDSC-F. Gene set enrichment analysis revealed that downregulated genes were enriched for organic molecule transport, whereas upregulated genes were associated with cuticle-related processes, many of which are expressed in muscle tissue (Figure 3C–D; Figure S2; Tables S5–S6). Considering that LDHA deficiency in humans is associated with skin abnormalities and exercise intolerance (Serrano-Lorenzo et al., 2022, Ozen, 2007), these gene expression changes are consistent with alterations in tissue properties relevant to organismal performance rather than increased metabolic disruption under lethal conditions. Together, these data further demonstrate that differences in nutrient composition or global metabolic state do not underlie the diet-dependent lethality of *Ldh* mutants and instead suggest that some other property of the feeding environment plays a central role in determining viability.

### *Ldh* mutant larvae are exercise intolerant and display locomotor defects

Having addressed differences in nutrient composition and global metabolic state, we next tested whether the diet-dependent lethality of *Ldh* mutants arises from the physical demands of feeding. Given that *Ldh* mutants show enrichment of gene expression changes in muscle-associated pathways, and that LDHA deficiency in humans is characterized by exercise intolerance, we hypothesized that differences in larval viability result from the amount of physical exertion required to consume each food type. Larvae feed at the interface between solid molasses agar and yeast paste, whereas they must burrow into semi-solid BDSC-F, a behavior likely requiring increased physical effort.

To test this hypothesis, we created a thin smear of BDSC-F on filter paper dampened with PBS (Figure S1A, see methods), thereby eliminating the need for larvae to burrow and mimicking the surface-feeding environment of M+Y plates. Strikingly, *Ldh* mutant larvae raised on this BDSC-F smear exhibited significantly higher survival than those reared within a cube of solid BDSC-F (Figure 4A), indicating that differences in viability are driven by the physical properties of the feeding environment. To confirm that these phenotypes arise specifically from loss of *Ldh*, we demonstrated that expressing a previously described *Ldh-GFP* rescue transgene in the mutant background restored viability under these conditions (Figure 4A,B,D,E). In contrast, supplementing BDSC-F with individual components of M+Y plates (molasses, yeast, or phosphoric acid) or antioxidants failed to improve mutant survival (Figure S1B–C), further demonstrating that diet-dependent lethality is not caused by nutritional composition or oxidative stress.

**Figure 4.**
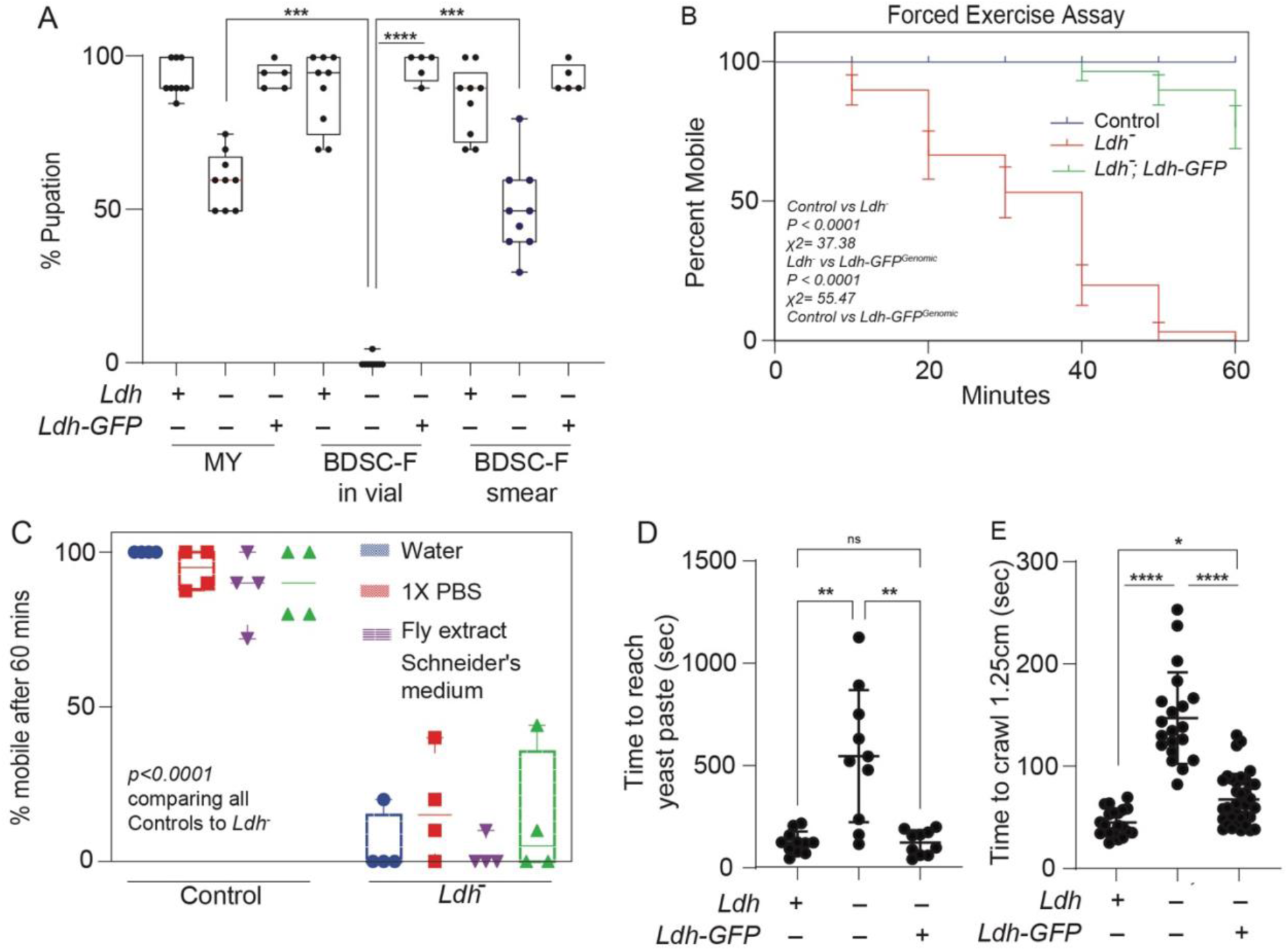
*Ldh* mutants exhibit exercise intolerance and impaired locomotor performance. (A) Percent pupation of control (*Ldh*^17^/+), *Ldh* mutant (*Ldh*^17^/*Ldh*^16^), and rescued *Ldh* mutant larvae expressing a previously described *Ldh*-GFP transgene in the *Ldh* mutant background. Larvae were raised on M+Y plates, BDSC-F in vials, or BDSC-F smeared on moist filter paper. Expression of the *Ldh*-GFP transgene restored viability under these conditions. Each experiment included ≥3 biological replicates with 10 larvae per replicate. Statistical analysis was performed using repeated-measures one-way ANOVA followed by a Holm–Šidák multiple-comparison test (\**P* < 0.0001). (B) Survival of larvae during a forced exercise assay. *Ldh* mutant larvae exhibited progressive mortality during 60 min of continuous forced movement, whereas control larvae survived throughout the assay and *Ldh* mutants expressing the *Ldh*-GFP rescue transgene only exhibited lethality after 40 min. Survival data were analyzed using a log-rank (Mantel–Cox) test (*N* = 30, *P* < 0.0001, χ² = 54.72). (C) Survival of control and *Ldh* mutant larvae during continuous swimming in water, 1× PBS, fly extract, or Schneider’s medium. *Ldh* mutants displayed significantly reduced survival in all media conditions. *N* = 4 biological replicates. Statistical analysis was performed using two-way ANOVA followed by a Holm–Šidák multiple-comparison test (*P* < 0.0001 for mutants versus controls in all conditions). (D) Time required for larvae to reach a yeast source. *Ldh* mutants were significantly delayed relative to controls, whereas expression of the *Ldh*-GFP rescue transgene in the *Ldh* mutant background restored performance to near-control levels. Statistical analysis was performed using a Mann–Whitney test (*N* = 10, *P* = 0.0005). (E) Crawling performance of control, *Ldh* mutant, and rescued *Ldh* mutant larvae expressing the *Ldh*-GFP transgene. *Ldh* mutants moved significantly more slowly than controls, and expression of the rescue transgene restored crawling performance. Statistical analysis was performed using a Mann–Whitney test (*N* = 20, *P* < 0.0001).

To directly test whether *Ldh* mutant larvae are exercise intolerant, we subjected animals to forced movement by gently prodding them with a paintbrush. Whereas controls and *Ldh* mutants expressing the rescuing transgene showed no lethality up to 40 minutes of stimulation, *Ldh* mutants began dying within 10 minutes, and all animals died by 50 minutes (Figure 4B). As an independent assay, we placed larvae in a small volume of liquid, which requires continuous swimming. More than 75% of control larvae survived for 60 minutes in water, PBS, fly extract, or Schneider’s media, whereas the majority of *Ldh* mutants died within the same period (Figures 4C and S5). Together, these results demonstrate that *Ldh* mutants exhibit pronounced exercise intolerance, closely mirroring the exertion-dependent phenotypes observed in individuals with LDHA deficiency.

Since loss of *Ldh* results in exercise intolerance, we next asked whether *Ldh* mutants also exhibit broader locomotor defects. To address this, we employed two complementary assays. First, we placed larvae at equal distances from a yeast source and measured the time required to reach the food. Second, we performed a standardized crawling assay (Post and Paululat, 2018). In both assays, *Ldh* mutants were significantly delayed in reaching the yeast (Figure 4D) and crawled more slowly than control or rescue (Figure 4E). Together, these results demonstrate that *Ldh* mutant larvae exhibit robust locomotor impairments.

Previous studies reported that RNAi-mediated depletion of *Ldh* disrupts muscle development, leading to structural abnormalities in larval muscles (Tixier et al., 2013). Because such defects could potentially explain the locomotor impairments observed in *Ldh* mutants, we examined whether muscle development was altered in our system. To do so, we dissected larval body-wall muscles from animals raised on M+Y, BDSC-F cubes, or BDSC-F smear preparations and stained them with phalloidin. In contrast to the earlier RNAi-based findings, we did not observe any gross defects in muscle morphology in *Ldh* loss-of-function mutants under any feeding condition (Figure S3A–F). Muscle architecture appeared comparable across genotypes and culture environments, indicating that loss of *Ldh* does not overtly disrupt muscle development in this context. This discrepancy may reflect differences in experimental conditions, the extent or timing of *Ldh* depletion, or distinctions between RNAi knockdown and the null alleles used here. Together, these results indicate that the locomotor and exercise-intolerance phenotypes of *Ldh* mutants are unlikely to result from gross defects in muscle development.

### Multiple tissues contribute to the diet-dependent *Ldh* mutant lethal phenotype

Although *Ldh* mutants exhibit normal muscle morphology, their locomotor and exercise-intolerance phenotypes suggest that muscle function may nonetheless be compromised. To assess whether restoring *Ldh* activity in muscle is sufficient to rescue viability, we expressed a previously described *UAS-Ldh* transgene using muscle-, neuronal-, and glia-specific drivers. However, *UAS-Ldh* expression in these cell types failed to rescue the lethality of *Ldh* mutants on BDSC-F (Figure S4), raising the possibility that *Ldh* activity is required in multiple tissues to support survival on this medium.

We next asked whether loss of *Ldh* within individual tissues contributes to the observed phenotypes. Starting with the muscle-specific GAL4 driver (*Mef2R-Gal4*) to express a previously described *UAS-Ldh*-RNAi transgene, we assessed the viability of larvae raised on BSDC-F and when placed in liquid media. Consistent with an essential role for Ldh in muscle, *Mef2R>Ldh-RNAi* resulted in approximately 40% lethality on solid BDSC-F (Figure 5A). Similarly, fewer than 50% of *Mef2R>Ldh-RNAi* larvae survived 60 minutes of continuous swimming across four liquid media conditions, whereas all control larvae survived (Figure 5B). Together, these findings indicate that loss of *Ldh* in muscle contributes substantially to the locomotor and survival defects observed in whole-animal mutants.

**Figure 5.**
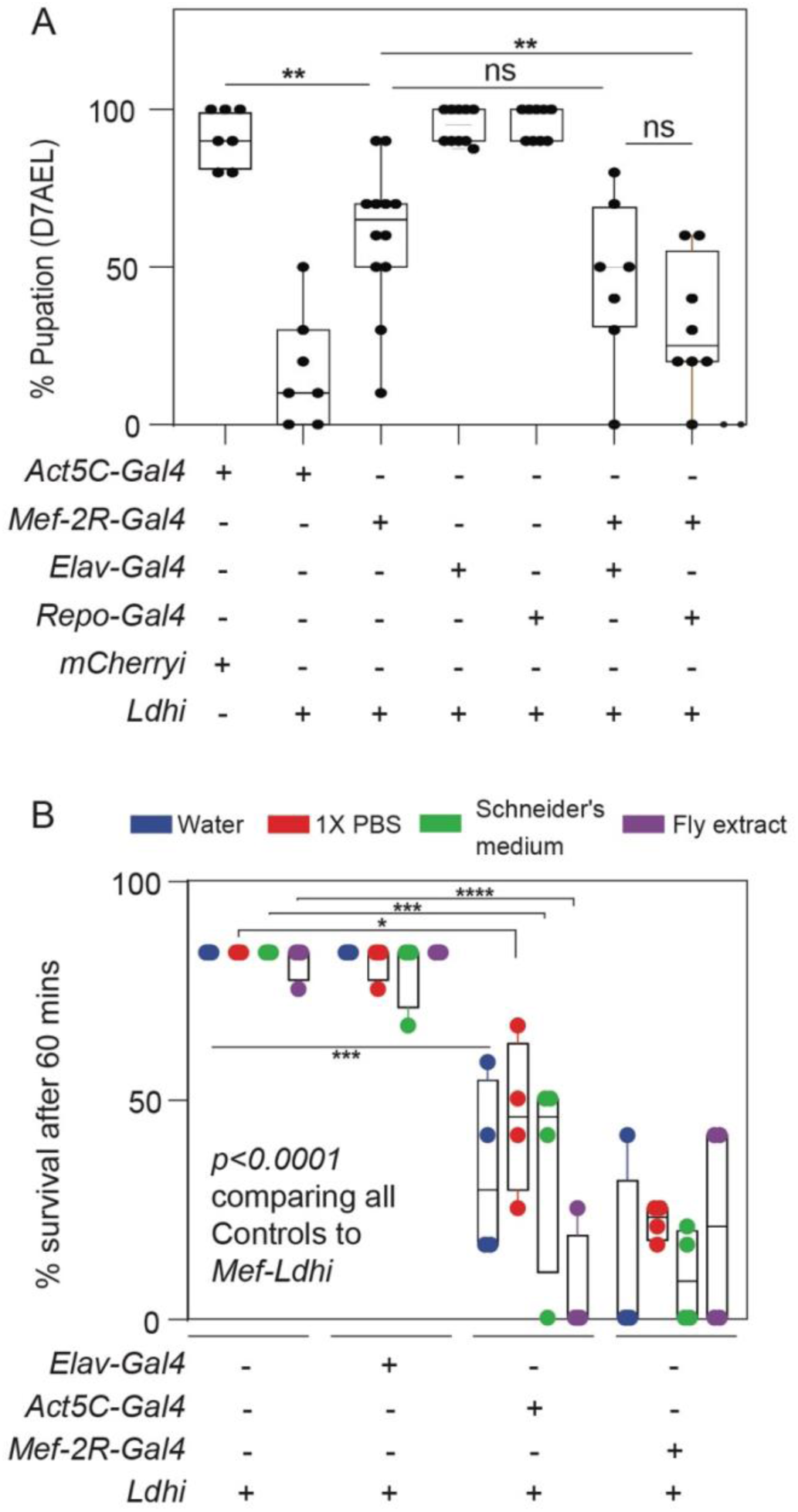
Multiple tissues contribute to the physiological requirements for *Ldh* during locomotor stress. (A) Percent pupation at 7 days AEL following tissue-specific knockdown of *Ldh* expression using *UAS-Ldh-RNAi*. RNAi was driven ubiquitously (*Act5C-GAL4*), in muscle (*Mef2R-GAL4*), neurons (*elav-GAL4*), or glia (*repo-GAL4*). *UAS-mCherry-RNAi* served as a control. Larvae were raised on BDSC-F in vials. Simultaneous RNAi knockdown of *Ldh* expression in muscle and neurons, as well as in muscle and glia, caused enhanced lethality relative to knockdown in a single tissue. *N* > 5 biological replicates. Statistical analysis was performed using two-way ANOVA followed by a Holm–Šidák multiple-comparison test (\**P* < 0.01). (B) Survival of larvae expressing *Ldh-RNAi* ubiquitously, in muscle, or in neurons during continuous swimming in water (blue), 1× PBS (red), fly extract (purple), or Schneider’s medium (green). Knockdown of *Ldh* significantly reduced survival relative to controls across all media conditions. *N* = 4 biological replicates. Statistical analysis was performed using two-way ANOVA followed by a Holm–Šidák multiple-comparison test (*P* < 0.0001 for all comparisons).

Because larval locomotion depends on coordinated activity between muscle and the peripheral nervous system, we next tested whether *Ldh* function in neurons or glia also contributes to survival. Motor neurons innervate body-wall muscles and are supported by peripheral glia that provide structural and metabolic support (Ikeda and Koenig, 1988, Menon et al., 2013), and disruption of lactate metabolism in glia has been shown to impair motor function (Bloom et al., 2022), while defects in either muscle or the peripheral nervous system (PNS) can compromise larval locomotion (Kottmeier et al., 2020, Corty and Coutinho-Budd, 2023). Consistent with a multi-tissue requirement, combined knockdown of *Ldh* in muscle and glia increased lethality to ∼60%, whereas knockdown in muscle and neurons produced a more modest effect (∼50%; Figure 5B). Together, these results demonstrate that *Ldh* activity across multiple tissues is required to support physical performance under conditions of increased mechanical demand. Hence, the diet-dependent lethality of *Ldh* mutants reflects the combined impact of impaired function across several tissues rather than a defect confined to muscle alone.

## DISCUSSION

Lactate dehydrogenase (LDH) plays a central role in cellular metabolism by coupling glycolytic flux to redox balance (Brooks, 2009, Brooks et al., 2022, Gladden, 2004, Van Voorhies, 2009, Li et al., 2019, Rai et al., 2025), yet the physiological consequences of disrupting this activity at the organismal level remain incompletely understood. Here, we demonstrate that *Drosophila Ldh* mutants exhibit a context-dependent phenotype in which viability is not determined solely by nutrient composition, but instead by the physical properties of the feeding environment. Under conditions that impose increased mechanical demand, such as feeding on solid food or sustained locomotion, *Ldh* mutants display severe exercise intolerance, closely mirroring key features of human LDHA deficiency (GSD-XI) (Maekawa et al., 1990, Kanno et al., 1980, Serrano-Lorenzo et al., 2022). These findings establish *Drosophila* as a powerful in vivo model for understanding how metabolic perturbations give rise to stress-dependent disease phenotypes.

A central insight from this study is that the lethality of *Ldh* mutants cannot be explained by differences in nutrient composition or global metabolic state. Although both metabolomic and transcriptomic analyses revealed that environmental conditions strongly influence systemic physiology, neither dataset identified signatures that correlate with viability. Indeed, *Ldh* mutants exhibited more extensive metabolic and transcriptional alterations under permissive conditions (M+Y) than under lethal conditions (BDSC-F), indicating that the extent of global physiological disruption does not predict survival. Instead, our data identify food consistency as a key determinant of viability, as reducing the mechanical demands of feeding by presenting food as a surface smear was sufficient to rescue viability, while multiple independent assays demonstrated that *Ldh* mutants exhibit pronounced locomotor impairments and exercise intolerance. Together, these findings support a model in which loss of *Ldh* creates a latent physiological defect that is largely tolerated under baseline conditions but becomes limiting under conditions that require sustained physical activity.

This exertion-dependent phenotype closely parallels the clinical presentation of LDHA deficiency in humans, where symptoms such as muscle pain, weakness, and rhabdomyolysis are typically triggered by physical activity rather than evident at rest (Serrano-Lorenzo et al., 2022, Kanno et al., 1980). At a mechanistic level, these findings are consistent with a model in which loss of LDH limits the regeneration of NAD⁺ during periods of high glycolytic demand, thereby constraining ATP production in metabolically active tissues such as muscle. Under conditions of increased exertion, this limitation would impair the ability to sustain movement, ultimately leading to organismal failure. Although this model remains to be tested directly, it provides a clear framework for interpreting the observed phenotypes and for guiding future studies.

Our tissue-specific analyses further reveal that the physiological requirement for *Ldh* is distributed across multiple interacting systems. While muscle-specific knockdown of *Ldh* impairs locomotion and reduces survival, it is not sufficient to recapitulate the full lethality of null mutants, and muscle-specific rescue fails to restore viability. Instead, survival requires *Ldh* activity across multiple tissues, including components of the peripheral nervous system. These findings are consistent with the concept that metabolic homeostasis during periods of high demand depends on coordinated interactions between tissues, rather than cell-autonomous function alone (Gonzalez-Gutierrez et al., 2020, Scott et al., 2023, Yellen, 2018, Brooks, 2009). In this context, LDH may support a system-level metabolic network that enables sustained neuromuscular performance.

More broadly, our results highlight the importance of considering environmental and behavioral context when evaluating the physiological consequences of metabolic divergence. Many metabolic disorders, including LDHA deficiency (Kanno et al., 1980, Kanno et al., 1988), present with mild or variable symptoms that become clinically significant only under conditions of stress or exertion. Our findings illustrate how the physiological consequences of metabolic perturbation can depend critically on context, with latent vulnerabilities emerging only under specific environmental or behavioral conditions. These observations have important implications for therapeutic strategies targeting metabolic enzymes such as LDHA. Specifically, they suggest that the efficacy and toxicity of LDHA-directed therapies may depend strongly on physiological context, with both therapeutic benefit and adverse effects becoming more pronounced during periods of elevated metabolic demand. Identifying the environmental, behavioral, and disease-associated conditions that place the greatest burden on LDH-dependent pathways may therefore help predict treatment-associated toxicities and identify disease states that are particularly dependent on LDH activity, thereby informing patient stratification and improving the therapeutic efficacy of LDHA-targeted interventions.

In summary, this study demonstrates that *Ldh* supports organismal resilience under conditions of increased physical demand, and that its loss leads to a conserved, stress-dependent phenotype mirroring human disease. By linking metabolic function to behaviorally relevant environmental challenges, this investigation establishes a framework for understanding how disruptions in core metabolic pathways translate into organismal pathology.

## MATERIALS AND METHODS

### *Drosophila melanogaster* husbandry and genetic analysis

Fly stocks were maintained at 25°C on Bloomington *Drosophila* Stock Center (BDSC) food. *Ldh* mutations were maintained in trans to the balancer *TM3, p{Dfd-GMR-nvYFP}, Sb^1^* (RRID: BDSC_23231). Unless noted, *Ldh* mutant larvae harbored a trans-heterozygous combination of *Ldh^16^* (RRID: BDSC_94698) and *Ldh^17^* (RRID: BDSC_94699) (Li et al., 2017). Muscle-, neuron-, and glia-specific knockdown of Ldh was conducted using transgenes that express GAL4 in the muscle *(P{w[+mC]=GAL4-Mef2.R}3*; RRID: BDSC_27390), neuron (*P{w[+mC]=GAL4-elav.L}2/CyO;* RRID: 8765) and glia (*P{w[+mC]=repo-Gal4.L Oatp30B[repoF3];* RRID: 602908). RNAi experiments were conducted using transgenes that targeted *Ldh* expression (RRID: BDSC_33640). The efficacy of the UAS-*Ldh*-RNAi transgene and Mef2-GAL4 driver has been validated previously (Li et al., 2017, Rai et al., 2025). Tissue-specific rescue experiments were performed using a UAS-*Ldh* transgene obtained from the Zurich ORFeome Project (Bischof et al., 2013), which was expressed in muscle, neurons, or glia using the tissue-specific GAL4 drivers described above. The previously described *Ldh-GFP^genomic^* rescue construct (RRID:BDSC_94704) was used as a positive control for the rescue experiments (Rai et al., 2024). FlyBase was used as a reference resource for gene, allele, and stock information throughout this study (Gramates et al., 2022, Ozturk-Colak et al., 2024).

### Larval feeding and viability assessment

Larvae were raised and staged as previously described (Li et al., 2017). Briefly, 50 virgin females and 25 males were placed in a mating bottle, and embryos were collected for 4 h on 35-mm molasses agar plates supplemented with a smear of yeast paste. Embryo collection plates were placed inside 60-mm Petri dishes and maintained at 25°C. At 24 h after egg laying (AEL), hatched larvae were transferred to one of four feeding conditions: (1) molasses agar plates supplemented with yeast paste (M+Y), (2) standard Bloomington *Drosophila* Stock Center food in vials (BDSC-F), (3) BDSC-F smear preparations, or (4) BDSC-F cubes (Figure S1A). The BDSC-F smear condition consisted of a thin layer of BDSC food spread across PBS-soaked filter paper in a 35-mm dish, thus allowing larvae to feed without a need to burrow into the food. In contrast, the BDSC-F cube condition consisted of a cube of BDSC food placed on filter paper in a 35-mm dish and surrounded by 100 μL of 1× PBS (Figure S1A). Larvae were maintained at 25°C, and viability was assessed by scoring pupation at 7 days AEL. Percent pupation was calculated using GraphPad Prism v10.1.2.

To test whether specific components of the M+Y diet could rescue *Ldh* mutant viability, BDSC-F was supplemented with either molasses (115 mL/L), yeast paste, or phosphoric acid (25 mL/L). Viability was assessed as described above.

### Sample collection and extraction for Metabolomics Analysis

*Ldh* mutant (*Ldh*^17^/*Ldh*^16^) and heterozygous control (*Ldh*^17^/+) eggs were collected on M+Y food. At 40–44 h after egg laying (AEL), 50 larvae were transferred to each of two M+Y plates and one BDSC-F cube preparation and maintained at 25°C. Fourteen hours later (∼54 h AEL), one of the M+Y plates was exposed to hypoxia (1% O_2_) for 6 h. Following hypoxia treatment, 25 mid-second-instar larvae (∼60 h AEL) were collected from each condition (M+Y, BDSC-F cubes, and M+Y under hypoxia) and processed for metabolomic analysis as previously described (Li and Tennessen, 2018).

Briefly, larvae were collected into prechilled 1.5 mL microcentrifuge tubes on ice, washed with ice-cold water, and immediately frozen in liquid nitrogen. Samples were transferred to pre-weighed 1.4 mm bead tubes, weighed using a Mettler Toledo XS105 balance, returned to liquid nitrogen, and stored at −80°C until extraction. Metabolites were extracted by homogenizing samples in 0.8 mL of prechilled (−20°C) 90% methanol containing 2 μg/mL succinic-d4 acid for 30 s at 6.45 m/s using a bead mill homogenizer in a 4°C temperature-controlled room. Homogenates were incubated at −20°C for 1 h and centrifuged at 20,000 × *g* for 5 min at 4°C. The resulting supernatants were submitted to the University of Colorado Anschutz Medical Campus for metabolomic analysis as previously described (Nemkov et al., 2019).

### Statistical analysis of metabolite data

Metabolomic data were analyzed using MetaboAnalyst 6.0 (Pang et al., 2024). Metabolite abundances were normalized to sample mass, log-transformed, and Pareto scaled prior to analysis. Relative metabolite abundances were plotted using GraphPad Prism v10.1.2. Statistical comparisons were performed using two-way ANOVA followed by Holm–Šidák multiple-comparison tests.

### RNAseq Analysis

Control (*Ldh*^17^/+) and *Ldh* mutant (*Ldh*^17^/*Ldh*^16^) eggs were collected on M+Y food. At 40–44 h after egg laying (AEL), 50 larvae were transferred to either M+Y plates or BDSC-F cube preparations and maintained at 25°C. Three biological replicates, each consisting of 25 mid-second-instar larvae (∼60 h AEL), were collected from each condition and washed twice with ice-cold 1× PBS. Total RNA was extracted from homogenized larvae using an RNeasy Kit (Qiagen; 74104). Library preparation and sequencing were performed at the Indiana University Center for Genomics and Bioinformatics.

RNA-seq reads were quantified using the Salmon pseudoaligner (Patro et al., 2017) against the *D. melanogaster* BDGP6.32 transcriptome, with the soft-masked genome assembly as decoy. Transcript-level abundance estimates were imported into DESeq2 using tximport for normalization and differential expression analysis (Soneson et al., 2015, Love et al., 2014). Log_2_ fold-change estimates were generated using lfcShrink, and genes with an adjusted *P*-value ≤ 0.05 and an absolute log_2_ fold change ≥ 1 were considered significantly differentially expressed. Venn diagrams were generated using the Matplotlib library in Python.

### GO analysis using PANGEA

RNAseq data were analyzed using PANGEA (Hu et al., 2023). Genes that were significantly upregulated or downregulated were analyzed for GO Enrichment using the GO Biological Process (BP), GO Molecular functions (MF) and preferred tissue (modEncode RNA_seq) sets (Gene Ontology et al., 2023).

### Immunofluorescence Assay

Body-wall muscles were dissected from early third-instar larvae (80 h AEL) in 1× phosphate-buffered saline (PBS; pH 7.0) and fixed in 4% paraformaldehyde in PBS for 30 min at room temperature. Samples were washed once with 1× PBS and twice with 0.3% PBT (PBS containing 0.3% Triton X-100) for 10 min per wash. Tissues were then blocked in 3% BSA prepared in 0.3% PBT for 30 min at room temperature and stained overnight at 4°C with Alexa Fluor 488-conjugated phalloidin (Molecular Probes). Following staining, samples were washed with 0.3% PBT, rinsed with 1× PBS, and mounted in VECTASHIELD mounting medium (Vector Laboratories; H-1200-10).

Muscle fillets were imaged using a Leica SP8 confocal microscope at the Indiana University Light Microscopy Imaging Center. Images were processed using FIJI, and figures were assembled in Adobe Illustrator 2020.

### Larval exercise assay

Larvae were raised on M+Y food at 25°C, and early third-instar larvae were selected for analysis. Groups of 10 larvae were aligned on a molasses agar plate lacking yeast paste and continuously stimulated by gently moving each larva back and forth with a paintbrush for 60 min. Survival was assessed throughout the assay based on larval responsiveness to tactile stimulation with the brush. Three biological replicates, each consisting of 10 larvae, were analyzed for both control (*Ldh*^17^/+) and *Ldh* mutant (*Ldh*^17^/*Ldh*^16^) animals. The survival time of each larva was recorded and analyzed using GraphPad Prism v10.1.2.

### Larval survival assay in liquid media

Larvae were raised on M+Y food at 25°C. Early third-instar larvae were transferred to a well of a 9-well glass spot test plate containing 200 μL of liquid medium (water, 1× PBS, Schneider’s medium, or fly extract). Note that fly extract was obtained from the *Drosophila* Genome Resource Center (Luhur et al., 2020). Survival was assessed every 2 min by gently prodding larvae with a paintbrush. For each condition, a minimum of 30 larvae were tested for control (*Ldh*^17^/+), *Ldh* mutant (*Ldh*^17^/*Ldh*^16^), and rescue strain (*Ldh*^17^/*Ldh*^16^*; Ldh-GFP^genomic^*). Data were plotted and analyzed using GraphPad Prism v10.1.2.

### Assay for larval attraction to yeast paste

Larvae were reared on M+Y food at 25°C. Early third-instar larvae were placed at one end of a molasses agar plate, while a measured aliquot of yeast paste was positioned at the opposite end. Larval movement was recorded for 1 h using a digital camera. Times were plotted and statistically analyzed using GraphPad Prism v10.1.2.

### Crawling assay

The crawling assay was adapted from a previous study (Post and Paululat, 2018). Early third-instar larvae raised on M+Y food were placed on a 60-mm plate lined with moist paper marked with a grid. Larvae were observed under a stereomicroscope, and the time required to travel 1.25 cm was recorded. Distance was determined by measuring the time required to traverse five consecutive 0.25-cm grid squares. Data were plotted and analyzed using GraphPad Prism v10.1.2.

## Data Availability

All strains and reagents are available upon request. All targeted metabolomics data described herein are included in Tables S1. RNA-seq data available in NCBI Gene Expression Omnibus (GEO; GSE337893).

## ACKNOWLEDGEMENTS

We thank the Bloomington Drosophila Stock Center (NIH P40OD018537) and FlyORF for providing fly stocks, the Drosophila Genomics Resource Center (NIH 2P40OD010949) for genomic reagents, Flybase (NIH 5U41HG000739), the IU Center for Genomics and Bioinformatics, and the Indiana University Light Microscopy Imaging Center. Targeted GC-MS analysis was conducted using instruments housed in the Indiana University Mass Spectrometry Facility, which was supported, in part, by NSF 1726633 to J.A.K.. and J.M.T. Metabolomic studies conducted by T.N. and A.D. were supported by the National Institute of Diabetes and Digestive and Kidney Diseases of the National Institutes of Health under Award Number R01DK136945. M.K. was supported by the Karen Bush and Daniel J. Watts Scholarship in Biotechnology and the Lawrence M. Blatt Biotechnology Internship Scholarship from Indiana University. S.A.S. was supported by the John R. and Wendy L. Kindig Fellowship, Robert Briggs Fellowship, Dona G. Graam Fellowship through Indiana University, and an American Heart Association Predoctoral Fellowship (25PRE1372770; https://doi.org/10.58275/AHA.25PRE1372770.pc.gr.227196). All other aspects of the research supported in this publication was supported by the NIGMS of the National Institute of Health under award R35GM119557 to J.M.T.

## SUPPLEMENTAL MATERIAL

**Table S1.** Metabolomic analysis of *Ldh*^17^/*Ldh*^16^ mutants (Ldh-_*) in comparison to *Ldh*^17^/+ heterozygous controls (HC_*), when fed with M+Y (*_MY), BDSC-F cubes (*_BF) and M+Y+ hypoxia (*_Hypoxia).

**Table S2.** List of significantly altered metabolites in *Ldh*^17^/*Ldh*^16^ mutants vs *Ldh*^17^/+ heterozygous controls, when fed with M+Y diet. Cutoff for significance is an adjusted *p-*value of <0.1 and a log2(FC) of ±1. Data generated using Metaboanalyst 6.0.

**Table S3.** List of significantly altered metabolites in *Ldh*^17^/*Ldh*^16^ mutants vs *Ldh*^17^/+ heterozygous controls, when fed with BDSC-Cubes. Cutoff for significance is an adjusted *p-*value of <0.1 and a log2(FC) of ±1. Data generated using Metaboanalyst 6.0.

**Table S4.** List of significantly altered genes in *Ldh*^17^/*Ldh*^16^ mutants (mut_*) vs *Ldh*^17^/+ heterozygous controls (ctrl_*), when fed with M+Y (*_MY) or BDSC-F Cubes (*_BDSC).

**Table S5.** Gene set enrichment analysis and Preferred tissue (modEncode RNA_seq) analysis for **downregulated** genes in *Ldh*^17^/*Ldh*^16^ mutants vs *Ldh*^17^/+ heterozygous control larvae fed with BDSC-F cubes cubes using PANGEA. Genes that were downregulated in both BDSC-F and M+Y were excluded from the analysis. Preferred tissue rows are marked with red.

**Table S6.** Gene set enrichment analysis and Preferred tissue (modEncode RNA_seq) analysis for upregulated genes in *Ldh*^17^/*Ldh*^16^ mutants vs *Ldh*^17^/+ heterozygous control larvae fed with BDSC-F cubes using PANGEA. Genes that were upregulated in both BDSC-F and M+Y were excluded from the analysis. Preferred tissue rows are marked with red.

## SUPPLEMENTAL FIGURE LEGENDS

**Figure S1. Addition of M+Y components to BDSC-F does not rescue Ldh mutant larval lethality.** (A) Images of BDSC-F in a vial, M+Y plate, BDSC-F cubes and BDSC-F smear, used for feeding larvae (left to right). (B) Percent pupation for control and Ldh mutants fed with BDSC-F in a vial supplemented with added water (control), molasses, yeast or phosphoric acid (PA). Each experiment had ≥ 3 replicates. Each replicate had 10 larvae. Statistical analysis was conducted using a two-way ANOVA followed by a Holm-Sidak test. ****P<0.0001

**Figure S2. Downregulated gene ontology for *Ldh* mutants in comparison to controls, when fed with BDSC-F cubes.** RNA-seq was used to analyze gene expression in L2 larvae of *Ldh* mutants relative to controls. (A) Genes exclusively downregulated in *Ldh* mutants compared to controls, when fed with BDSC-F cubes, were analyzed for gene enrichment based on GO- Biological Processes and GO-Molecular functions, using PANGEA. (B) Plot showing the tissue-enrichment of downregulated genes. Three replicates were used for the transcriptomic experiment with each replicate having 20 larvae.

**Figure S3. Larval muscles remain morphologically intact in *Ldh* mutants in comparison to controls, independent of diet.** Representative confocal images of dissected muscle fillets from control (A-C) and *Ldh* mutants (D-F) larvae fed with M+Y, BDSC-F Cubes and BDSC-F Smear. Muscles are shown by Phalloidin staining, shown in green. Scale bar = 50um from A, applies to B-F.

**Figure S4. Expression of Ldh in the muscles, neurons or glia does not rescue *Ldh* mutant lethality on BDSC-F.** Graph showing the percent pupation at 7 days after egg-laying (D7AEL) for larvae raised in vials of BDSC-F. *UAS-Ldh* was expressed in the background of *Ldh* mutants either in the muscles (using *Mef-2R-gal4*), neurons (using *Elav-Gal4)* or in the glia (using *Repo-Gal4). N=4*. Each replicate had 10 larvae. Statistical analysis was conducted using a RM one-way ANOVA followed by a Holm-Sidak test. *\*\*\*P<0.0001*.

**Figure S5.** Survival of *Ldh* mutant larvae expressing a *Ldh-GFP^genomic^* rescue construct during continuous swimming in water, 1× PBS, fly extract, or Schneider’s medium. *N* = 4 biological replicates. Error bars represent standard deviation.

## References

Abu-Shumays, R. L. & Fristrom, J. W. 1997. Imp-L3, A 20-hydroxyecdysone-responsive gene encodes Drosophila lactate dehydrogenase: structural characterization and developmental studies. Dev Genet, 20, 11–22.

Bartoloni, B., Mannelli, M., Gamberi, T. & Fiaschi, T. 2024. The Multiple Roles of Lactate in the Skeletal Muscle. Cells, 13.

Bischof, J., Bjorklund, M., Furger, E., Schertel, C., Taipale, J. & Basler, K. 2013. A versatile platform for creating a comprehensive Uas-ORFeome library in Drosophila. Development, 140, 2434–42.

Bloom, A. J., Hackett, A. R., Strickland, A., Yamada, Y., Ippolito, J., Schmidt, R. E., Sasaki, Y., Diantonio, A. & Milbrandt, J. 2022. Disruption of lactate metabolism in the peripheral nervous system leads to motor-selective deficits. bioRxiv, 2022.06.29.497865.

Brand, A., Singer, K., Koehl, G. E., Kolitzus, M., Schoenhammer, G., Thiel, A., Matos, C., Bruss, C., Klobuch, S., Peter, K., Kastenberger, M., Bogdan, C., Schleicher, U., Mackensen, A., Ullrich, E., Fichtner-Feigl, S., Kesselring, R., Mack, M., Ritter, U., Schmid, M., Blank, C., Dettmer, K., Oefner, P. J., Hoffmann, P., Walenta, S., Geissler, E. K., Pouyssegur, J., Villunger, A., Steven, A., Seliger, B., Schreml, S., Haferkamp, S., Kohl, E., Karrer, S., Berneburg, M., Herr, W., Mueller-Klieser, W., Renner, K. & Kreutz, M. 2016. Ldha-Associated Lactic Acid Production Blunts Tumor Immunosurveillance by T and Nk Cells. Cell Metab, 24, 657–671.

Brooks, G. A. 2009. Cell-cell and intracellular lactate shuttles. J Physiol, 587, 5591–600.

Brooks, G. A. 2018. The Science and Translation of Lactate Shuttle Theory. Cell Metab, 27, 757–785.

Brooks, G. A., Arevalo, J. A., Osmond, A. D., Leija, R. G., Curl, C. C. & Tovar, A. P. 2022. Lactate in contemporary biology: a phoenix risen. J Physiol, 600, 1229–1251.

Chen, X., Liu, L., Kang, S., Gnanaprakasam, J. R. & Wang, R. 2023. The lactate dehydrogenase (Ldh) isoenzyme spectrum enables optimally controlling T cell glycolysis and differentiation. Sci Adv, 9, eadd9554.

Corty, M. M. & Coutinho-Budd, J. 2023. Drosophila glia take shape to sculpt the nervous system. Curr Opin Neurobiol, 79, 102689.

Dai, M., Wang, L., Yang, J., Chen, J., Dou, X., Chen, R., Ge, Y. & Lin, Y. 2023. Ldha as a regulator of T cell fate and its mechanisms in disease. Biomed Pharmacother, 158, 114164.

Fang, Y., Li, Z., Yang, L., Li, W., Wang, Y., Kong, Z., Miao, J., Chen, Y., Bian, Y. & Zeng, L. 2024. Emerging roles of lactate in acute and chronic inflammation. Cell Commun Signal, 22, 276.

Farhana, A. & Lappin, S. L. 2026. Biochemistry, Lactate Dehydrogenase. *StatPearls*. Treasure Island (Fl).

Ferguson, B. S., Rogatzki, M. J., Goodwin, M. L., Kane, D. A., Rightmire, Z. & Gladden, L. B. 2018. Lactate metabolism: historical context, prior misinterpretations, and current understanding. Eur J Appl Physiol, 118, 691–728.

Frame, A. K., Robinson, J. W., Mahmoudzadeh, N. H., Tennessen, J. M., Simon, A. F. & Cumming, R. C. 2023. Aging and memory are altered by genetically manipulating lactate dehydrogenase in the neurons or glia of flies. Aging (Albany Ny), 15, 947–981.

Gene Ontology, C., Aleksander, S. A., Balhoff, J., Carbon, S., Cherry, J. M., Drabkin, H. J., Ebert, D., Feuermann, M., Gaudet, P., Harris, N. L., Hill, D. P., Lee, R., Mi, H., Moxon, S., Mungall, C. J., Muruganugan, A., Mushayahama, T., Sternberg, P. W., Thomas, P. D., Van Auken, K., Ramsey, J., Siegele, D. A., Chisholm, R. L., Fey, P., Aspromonte, M. C., Nugnes, M. V., Quaglia, F., Tosatto, S., Giglio, M., Nadendla, S., Antonazzo, G., Attrill, H., Dos Santos, G., Marygold, S., Strelets, V., Tabone, C. J., Thurmond, J., Zhou, P., Ahmed, S. H., Asanitthong, P., Luna Buitrago, D., Erdol, M. N., Gage, M. C., Ali Kadhum, M., Li, K. Y. C., Long, M., Michalak, A., Pesala, A., Pritazahra, A., Saverimuttu, S. C. C., Su, R., Thurlow, K. E., Lovering, R. C., Logie, C., Oliferenko, S., Blake, J., Christie, K., Corbani, L., Dolan, M. E., Drabkin, H. J., Hill, D. P., Ni, L., Sitnikov, D., Smith, C., Cuzick, A., Seager, J., Cooper, L., Elser, J., Jaiswal, P., Gupta, P., Jaiswal, P., Naithani, S., Lera-Ramirez, M., Rutherford, K., Wood, V., De Pons, J. L., Dwinell, M. R., Hayman, G. T., Kaldunski, M. L., Kwitek, A. E., Laulederkind, S. J. F., Tutaj, M. A., Vedi, M., Wang, S. J., D’eustachio, P., Aimo, L., Axelsen, K., Bridge, A., Hyka-Nouspikel, N., Morgat, A., Aleksander, S. A., Cherry, J. M., Engel, S. R., Karra, K., Miyasato, S. R., Nash, R. S., Skrzypek, M. S., Weng, S., Wong, E. D., Bakker, E., et al. 2023. The Gene Ontology knowledgebase in 2023. Genetics, 224.

Gladden, L. B. 2004. Lactate metabolism: a new paradigm for the third millennium. J Physiol, 558, 5–30.

Gonzalez-Gutierrez, A., Ibacache, A., Esparza, A., Barros, L. F. & Sierralta, J. 2020. Neuronal lactate levels depend on glia-derived lactate during high brain activity in Drosophila. Glia, 68, 1213–1227.

Gramates, L. S., Agapite, J., Attrill, H., Calvi, B. R., Crosby, M. A., Dos Santos, G., Goodman, J. L., Goutte-Gattat, D., Jenkins, V. K., Kaufman, T., Larkin, A., Matthews, B. B., Millburn, G., Strelets, V. B. & The Flybase, C. 2022. FlyBase: a guided tour of highlighted features. Genetics, 220.

Herranz, H. & Cohen, S. M. 2017. Drosophila as a Model to Study the Link between Metabolism and Cancer. J Dev Biol, 5.

Hu, Y., Comjean, A., Attrill, H., Antonazzo, G., Thurmond, J., Chen, W., Li, F., Chao, T., Mohr, S. E., Brown, N. H. & Perrimon, N. 2023. Pangea: a new gene set enrichment tool for Drosophila and common research organisms. Nucleic Acids Res, 51, W419–W426.

Ikeda, K. & Koenig, J. H. 1988. Morphological identification of the motor neurons innervating the dorsal longitudinal flight muscle of Drosophila melanogaster. J Comp Neurol, 273, 436–44.

Kanno, T., Sudo, K., Maekawa, M., Nishimura, Y., Ukita, M. & Fukutake, K. 1988. Lactate dehydrogenase M-subunit deficiency: a new type of hereditary exertional myopathy. Clin Chim Acta, 173, 89–98.

Kanno, T., Sudo, K., Takeuchi, I., Kanda, S., Honda, N., Nishimura, Y. & Oyama, K. 1980. Hereditary deficiency of lactate dehydrogenase M-subunit. Clin Chim Acta, 108, 267–76.

Kim, Y., Dube, S. E. & Park, C. B. 2025. Brain energy homeostasis: the evolution of the astrocyte-neuron lactate shuttle hypothesis. Korean J Physiol Pharmacol, 29, 1–8.

Kottmeier, R., Bittern, J., Schoofs, A., Scheiwe, F., Matzat, T., Pankratz, M. & Klambt, C. 2020. Wrapping glia regulates neuronal signaling speed and precision in the peripheral nervous system of Drosophila. Nat Commun, 11, 4491.

Kumar, S., Sahu, N., Jawaid, T., Jayasingh Chellammal, H. S. & Upadhyay, P. 2025. Dual role of lactate in human health and disease. Front Physiol, 16, 1621358.

Li, H., Chawla, G., Hurlburt, A. J., Sterrett, M. C., Zaslaver, O., Cox, J., Karty, J. A., Rosebrock, A. P., Caudy, A. A. & Tennessen, J. M. 2017. Drosophila larvae synthesize the putative oncometabolite L-2-hydroxyglutarate during normal developmental growth. Proc Natl Acad Sci U S A, 114, 1353–1358.

Li, H., Rai, M., Buddika, K., Sterrett, M. C., Luhur, A., Mahmoudzadeh, N. H., Julick, C. R., Pletcher, R. C., Chawla, G., Gosney, C. J., Burton, A. K., Karty, J. A., Montooth, K. L., Sokol, N. S. & Tennessen, J. M. 2019. Lactate dehydrogenase and glycerol-3-phosphate dehydrogenase cooperatively regulate growth and carbohydrate metabolism during Drosophila melanogaster larval development. Development, 146.

Li, H. & Tennessen, J. M. 2018. Preparation of Drosophila Larval Samples for Gas Chromatography-Mass Spectrometry (Gc-Ms)-based Metabolomics. J Vis Exp.

Liberti, M. V. & Locasale, J. W. 2020. Histone Lactylation: A New Role for Glucose Metabolism. Trends Biochem Sci, 45, 179–182.

Lin, Y., Wang, Y. & Li, P. F. 2022. Mutual regulation of lactate dehydrogenase and redox robustness. Front Physiol, 13, 1038421.

Liu, W., Wang, Y., Bozi, L. H. M., Fischer, P. D., Jedrychowski, M. P., Xiao, H., Wu, T., Darabedian, N., He, X., Mills, E. L., Burger, N., Shin, S., Reddy, A., Sprenger, H. G., Tran, N., Winther, S., Hinshaw, S. M., Shen, J., Seo, H. S., Song, K., Xu, A. Z., Sebastian, L., Zhao, J. J., Dhe-Paganon, S., Che, J., Gygi, S. P., Arthanari, H. & Chouchani, E. T. 2023. Lactate regulates cell cycle by remodelling the anaphase promoting complex. Nature, 616, 790–797.

Liu, W., Yang, R., Zhan, Y., Yang, X., Zeng, H., Chen, B., Zeng, J., Hu, T., Hu, J., Xiao, Q., Shao, Y. & Chen, X. 2025. Lactate and lactylation: emerging roles in autoimmune diseases and metabolic reprogramming. Front Immunol, 16, 1589853.

Llibre, A., Kucuk, S., Gope, A., Certo, M. & Mauro, C. 2025. Lactate: A key regulator of the immune response. Immunity, 58, 535–554.

Long, D. M., Frame, A. K., Reardon, P. N., Cumming, R. C., Hendrix, D. A., Kretzschmar, D. & Giebultowicz, J. M. 2020. Lactate dehydrogenase expression modulates longevity and neurodegeneration in Drosophila melanogaster. Aging (Albany Ny), 12, 10041–10058.

Love, M. I., Huber, W. & Anders, S. 2014. Moderated estimation of fold change and dispersion for Rna-seq data with DESeq2. Genome Biol, 15, 550.

Luhur, A., Mariyappa, D., Klueg, K. M., Buddika, K., Tennessen, J. M. & Zelhof, A. C. 2020. Adapting Drosophila melanogaster Cell Lines to Serum-Free Culture Conditions. G3 (Bethesda), 10, 4541–4551.

Ma, Z., Yang, J., Jia, W., Li, L., Li, Y., Hu, J., Luo, W., Li, R., Ye, D. & Lan, P. 2025. Histone lactylation-driven B7-H3 expression promotes tumor immune evasion. Theranostics, 15, 2338–2359.

Maekawa, M., Sudo, K., Kanno, T. & Li, S. S. 1990. Molecular characterization of genetic mutation in human lactate dehydrogenase-A (M) deficiency. Biochem Biophys Res Commun, 168, 677–82.

Menon, K. P., Carrillo, R. A. & Zinn, K. 2013. Development and plasticity of the Drosophila larval neuromuscular junction. Wiley Interdiscip Rev Dev Biol, 2, 647–70.

Merkle, S., Favor, J., Graw, J., Hornhardt, S. & Pretsch, W. 1992. Hereditary lactate dehydrogenase A-subunit deficiency as cause of early postimplantation death of homozygotes in Mus musculus. Genetics, 131, 413–21.

Nemkov, T., Reisz, J. A., Gehrke, S., Hansen, K. C. & D’alessandro, A. 2019. High-Throughput Metabolomics: Isocratic and Gradient Mass Spectrometry-Based Methods. Methods Mol Biol, 1978, 13–26.

Ozen, H. 2007. Glycogen storage diseases: new perspectives. World J Gastroenterol, 13, 2541–53.

Ozturk-Colak, A., Marygold, S. J., Antonazzo, G., Attrill, H., Goutte-Gattat, D., Jenkins, V. K., Matthews, B. B., Millburn, G., Dos Santos, G., Tabone, C. J. & Flybase, C. 2024. FlyBase: updates to the Drosophila genes and genomes database. Genetics, 227.

Pang, Z., Lu, Y., Zhou, G., Hui, F., Xu, L., Viau, C., Spigelman, A. F., Macdonald, P. E., Wishart, D. S., Li, S. & Xia, J. 2024. MetaboAnalyst 6.0: towards a unified platform for metabolomics data processing, analysis and interpretation. Nucleic Acids Res.

Park, Y. J., Lu, T. C., Jackson, T., Goodman, L. D., Ran, L., Chen, J., Liang, C. Y., Harrison, E., Ko, C., Chen, X., Wang, B., Hsu, A. L., Ochoa, E., Bieniek, K. F., Yamamoto, S., Zhu, Y., Zheng, H., Qi, Y., Bellen, H. J. & Li, H. 2025. Distinct systemic impacts of Aβ42 and Tau revealed by whole-organism snrna-seq. Neuron, 113, 2065–2082.e8.

Peng, M., Yin, N., Chhangawala, S., Xu, K., Leslie, C. S. & Li, M. O. 2016. Aerobic glycolysis promotes T helper 1 cell differentiation through an epigenetic mechanism. Science, 354, 481–484.

Post, Y. & Paululat, A. 2018. Muscle Function Assessment Using a Drosophila Larvae Crawling Assay. Bio Protoc, 8, e2933.

Rabinowitz, J. D. & Enerback, S. 2020. Lactate: the ugly duckling of energy metabolism. Nat Metab, 2, 566–571.

Rai, M., Carter, S. M., Shefali, S. A., Chawla, G. & Tennessen, J. M. 2024. Characterization of genetic and molecular tools for studying the endogenous expression of Lactate dehydrogenase in Drosophila melanogaster. PLos One, 19, e0287865.

Rai, M., Li, H., Policastro, R. A., Pepin, R., Zentner, G. E., Nemkov, T., D’alessandro, A. & Tennessen, J. M. 2025. Glycolytic disruption restricts Drosophila melanogaster larval growth via the cytokine Upd3. PLos Genet, 21, e1011690.

Rechsteiner, M. C. 1970. Drosophila lactate dehydrogenase and alpha-glycerolphosphate dehydrogenase: distribution and change in activity during development. J Insect Physiol, 16, 1179–92.

Sanchez, P. K. M., Khazaei, M., Gatineau, E., Geravandi, S., Lupse, B., Liu, H., Dringen, R., Wojtusciszyn, A., Gilon, P., Maedler, K. & Ardestani, A. 2021. Ldha is enriched in human islet alpha cells and upregulated in type 2 diabetes. Biochem Biophys Res Commun, 568, 158–166.

Scott, H., Novikov, B., Ugur, B., Allen, B., Mertsalov, I., Monagas-Valentin, P., Koff, M., Baas Robinson, S., Aoki, K., Veizaj, R., Lefeber, D. J., Tiemeyer, M., Bellen, H. & Panin, V. 2023. Glia-neuron coupling via a bipartite sialylation pathway promotes neural transmission and stress tolerance in Drosophila. Elife, 12.

Serrano-Lorenzo, P., Rabasa, M., Esteban, J., Hidalgo Mayoral, I., Dominguez-Gonzalez, C., Blanco-Echevarria, A., Garrido-Moraga, R., Lucia, A., Blazquez, A., Rubio, J. C., Palma-Milla, C., Arenas, J. & Martin, M. A. 2022. Clinical, Biochemical, and Molecular Characterization of Two Families with Novel Mutations in the Ldha Gene (Gsd Xi). Genes (Basel), 13.

Seth, P., Csizmadia, E., Hedblom, A., Vuerich, M., Xie, H., Li, M., Longhi, M. S. & Wegiel, B. 2017. Deletion of Lactate Dehydrogenase-A in Myeloid Cells Triggers Antitumor Immunity. Cancer Res, 77, 3632–3643.

Soneson, C., Love, M. I. & Robinson, M. D. 2015. Differential analyses for Rna-seq: transcript-level estimates improve gene-level inferences. F1000Res, 4, 1521.

Tendulkar, S., Wu, T., Strickland, A., Hackett, A. R., Sato-Yamada, Y., Mao, X., Sasaki, Y., Milbrandt, J., Bloom, A. J. & Diantonio, A. 2025. Dysregulated lactate metabolism synergizes with Als genetic risk factors to accelerate motor decline. bioRxiv.

Tendulkar, S., Wu, T., Strickland, A., Hackett, A. R., Sato-Yamada, Y., Mao, X., Sasaki, Y., Milbrandt, J., Bloom, A. J. & Diantonio, A. 2026. Dysregulated lactate metabolism synergizes with Als genetic risk factors to accelerate motor decline. PLos One, 21, e0347135.

Tennessen, J. M., Baker, K. D., Lam, G., Evans, J. & Thummel, C. S. 2011. The Drosophila estrogen-related receptor directs a metabolic switch that supports developmental growth. Cell Metab, 13, 139–48.

Tixier, V., Bataille, L., Etard, C., Jagla, T., Weger, M., Daponte, J. P., Strahle, U., Dickmeis, T. & Jagla, K. 2013. Glycolysis supports embryonic muscle growth by promoting myoblast fusion. Proc Natl Acad Sci U S A, 110, 18982–7.

Valvona, C. J., Fillmore, H. L., Nunn, P. B. & Pilkington, G. J. 2016. The Regulation and Function of Lactate Dehydrogenase A: Therapeutic Potential in Brain Tumor. Brain Pathol, 26, 3–17.

Van Voorhies, W. A. 2009. Metabolic function in Drosophila melanogaster in response to hypoxia and pure oxygen. J Exp Biol, 212, 3132–41.

Wang, C. W., Purkayastha, A., Jones, K. T., Thaker, S. K. & Banerjee, U. 2016. In vivo genetic dissection of tumor growth and the Warburg effect. Elife, 5.

Wen, H., Zhang, P., Zhao, J., Liu, Y., Wan, L., Li, H., Yi, J. & Li, X. 2025. Metabolic alterations driven by Ldha in CD8 + T cells promote immune evasion and therapy resistance in Nsclc. Sci Rep, 15, 24440.

Xiao, X., Zhang, J., Lang, Y., Cai, L., Yang, Q., Liu, K., Ji, S., Ju, X. & Liu, F. 2023. Associations of lactate dehydrogenase with risk of renal outcomes and cardiovascular mortality in individuals with diabetic kidney disease. Diabetes Res Clin Pract, 203, 110838.

Yao, S., Xu, M. D., Wang, Y., Zhao, S. T., Wang, J., Chen, G. F., Chen, W. B., Liu, J., Huang, G. B., Sun, W. J., Zhang, Y. Y., Hou, H. L., Li, L. & Sun, X. D. 2023. Astrocytic lactate dehydrogenase A regulates neuronal excitability and depressive-like behaviors through lactate homeostasis in mice. Nat Commun, 14, 729.

Yellen, G. 2018. Fueling thought: Management of glycolysis and oxidative phosphorylation in neuronal metabolism. J Cell Biol, 217, 2235–2246.

Zhang, D., Tang, Z., Huang, H., Zhou, G., Cui, C., Weng, Y., Liu, W., Kim, S., Lee, S., Perez-Neut, M., Ding, J., Czyz, D., Hu, R., Ye, Z., He, M., Zheng, Y. G., Shuman, H. A., Dai, L., Ren, B., Roeder, R. G., Becker, L. & Zhao, Y. 2019. Metabolic regulation of gene expression by histone lactylation. Nature, 574, 575–580.

Zhu, W., Ma, Y., Guo, W., Lu, J., Li, X., Wu, J., Qin, P., Zhu, C. & Zhang, Q. 2022. Serum Level of Lactate Dehydrogenase is Associated with Cardiovascular Disease Risk as Determined by the Framingham Risk Score and Arterial Stiffness in a Health-Examined Population in China. Int J Gen Med, 15, 11–17.

