## Supplemental Figures S1-S5 for "*Drosophila melanogaster* lactate dehydrogenase deficiency recapitulates the exercise intolerance of human glycogen storage disease type XI"

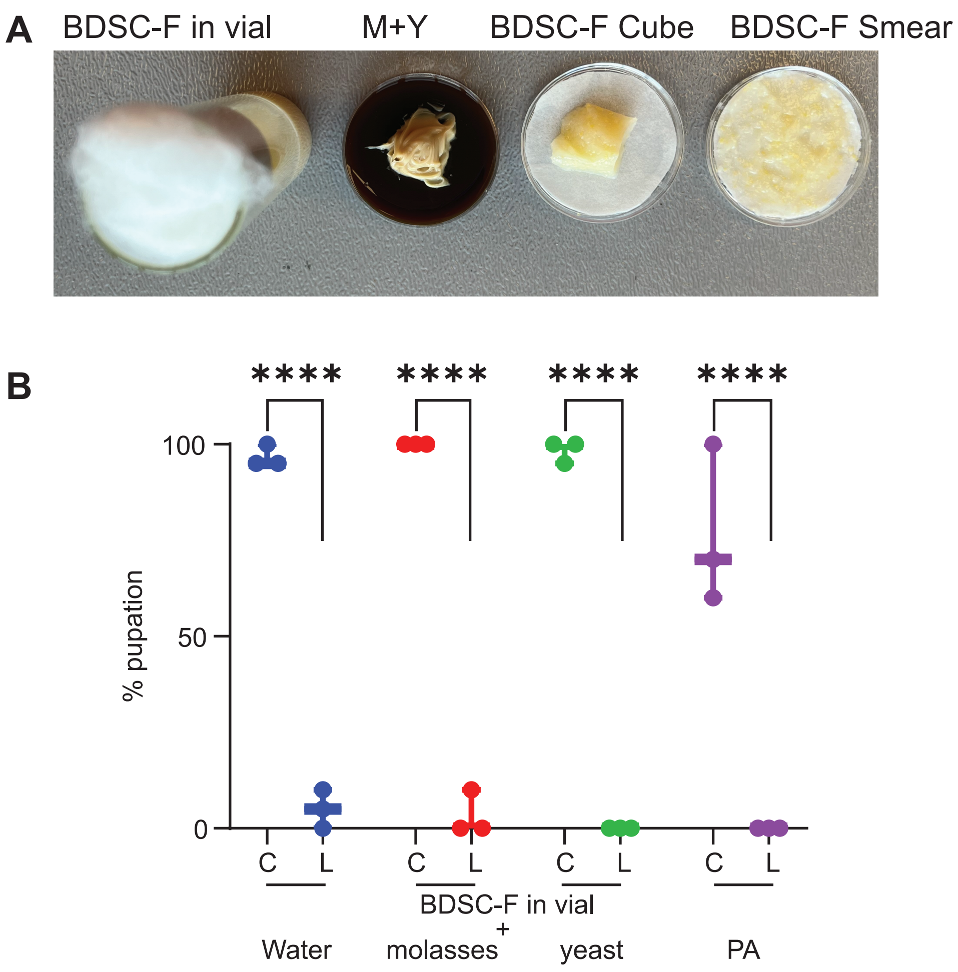


**Figure S1. Addition of M+Y components to BDSC-F does not rescue Ldh mutant larval lethality.** (A) Images of BDSC-F in a vial, M+Y plate, BDSC-F cubes and BDSC-F smear, used for feeding larvae (left to right). (B) Percent pupation for control and Ldh mutants fed with BDSC-F in a vial supplemented with added water (control), molasses, yeast or phosphoric acid (PA). Each experiment had ≥ 3 replicates. Each replicate had 10 larvae. Statistical analysis was conducted using a two-way ANOVA followed by a Holm-Sidak test. ****P<0.0001

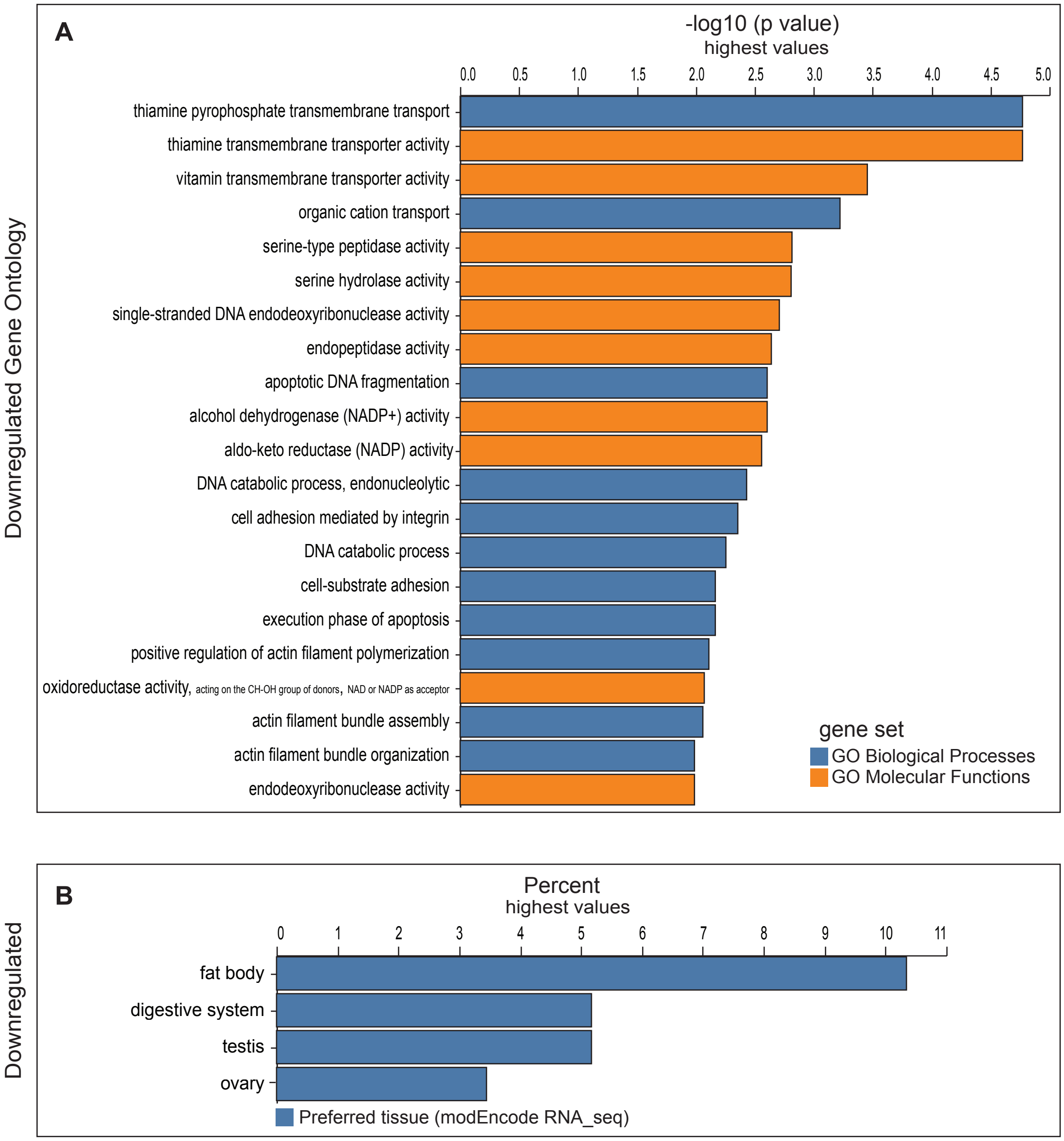


**Figure S2. Downregulated gene ontology for *Ldh* mutants in comparison to controls, when fed with BDSC-F cubes.** RNA-seq was used to analyze gene expression in L2 larvae of *Ldh* mutants relative to controls. (A) Genes exclusively downregulated in *Ldh* mutants compared to controls, when fed with BDSC-F cubes, were analyzed for gene enrichment based on GO- Biological Processes and GO- Molecular functions, using PANGEA. (B) Plot showing the tissue-enrichment of downregulated genes. Three replicates were used for the transcriptomic experiment with each replicate having 20 larvae.

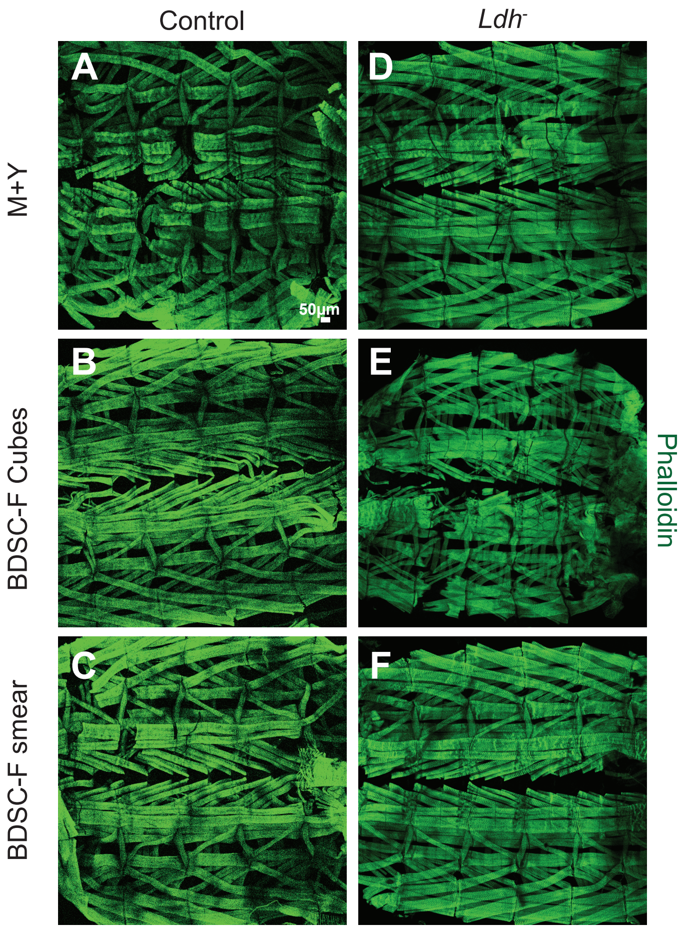


**Figure S3. Larval muscles remain morphologically intact in *Ldh* mutants in comparison to controls, independent of diet.** Representative confocal images of dissected muscle fillets from control (A-C) and *Ldh* mutants (D-F) larvae fed with M+Y, BDSC-F Cubes and BDSC-F Smear. Muscles are shown by Phalloidin staining, shown in green. Scale bar = 50um from A, applies to B-F.

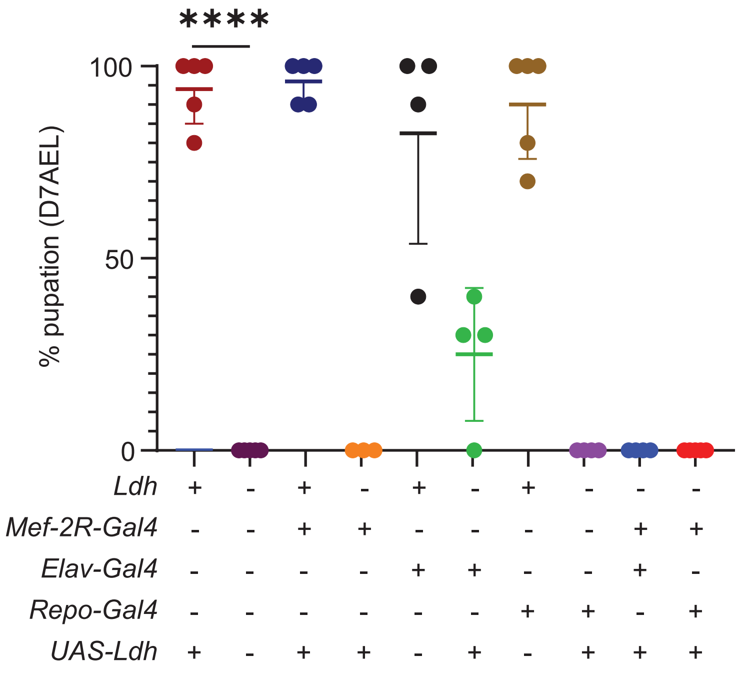


**Figure S4. Expression of Ldh in the muscles, neurons or glia does not rescue *Ldh* mutant lethality on BDSC-F.** Graph showing the percent pupation at 7 days after egg-laying (D7AEL) for larvae raised in vials of BDSC-F. *UAS-Ldh* was expressed in the background of *Ldh* mutants either in the muscles (using *Mef-2R-gal4*), neurons (using *Elav-Gal4)* or in the glia (using *Repo-Gal4). N=4*. Each replicate had 10 larvae. Statistical analysis was conducted using a RM one-way ANOVA followed by a Holm-Sidak test. ****P<0.0001.*

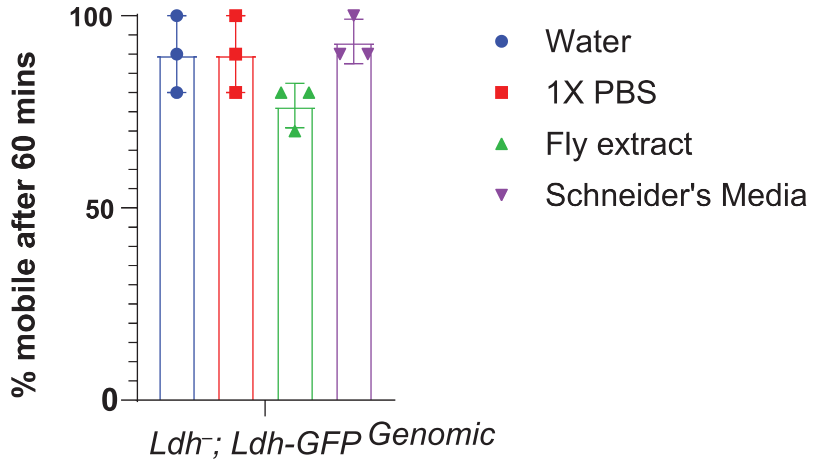


**Figure S5.** Survival of *Ldh* mutant larvae expressing a *Ldh-GFP*$^{genomic}$ rescue construct during continuous swimming in water, 1× PBS, fly extract, or Schneider's medium. *N* = 4 biological replicates. Error bars represent standard deviation.
